# RNA-binding proteins with mixed charge domains self-assemble and aggregate in Alzheimer’s Disease

**DOI:** 10.1101/243014

**Authors:** Isaac Bishof, Eric B. Dammer, Duc M. Duong, Marla Gearing, James J. Lah, Allan I. Levey, Nicholas T. Seyfried

**Affiliations:** Department of Biochemistry Emory University School of Medicine, Atlanta, GA, 30322; Department of Neurology Emory University School of Medicine, Atlanta, GA, 30322; Department of Pathology and Laboratory Medicine Emory University School of Medicine, Atlanta, GA, 30322; Center for Neurodegenerative Diseases, Emory University School of Medicine, Atlanta, GA, 30322

**Keywords:** protein-protein interaction, protein aggregation, systems biology, Tau protein, neurodegeneration, mass spectrometry, proteomics, RNA-binding protein, intrinsically disordered protein, RNA processing

## Abstract

U1 small nuclear ribonucleoprotein 70 kDa (U1-70K) and other RNA binding proteins (RBPs) are mislocalized to cytoplasmic neurofibrillary Tau aggregates in Alzheimer’s disease (AD), yet understanding of the mechanisms that cause their aggregation is limited. Many RBPs that aggregate in neurodegenerative diseases self-assemble into RNA granules through intrinsically disordered low complexity (LC) domains. We report here that a LC domain within U1-70K of mixed charge, containing highly repetitive complementary repeats of basic (R/K) and acidic (D/E) residues, shares many of the same properties of the Q/N-rich LC domains found in the RBPs TDP-43 and FUS. These properties include the ability to self-assemble into oligomers, and to form nuclear granules. To analyze the functional roles of the U1-70K LC domains, we performed co-immunoprecipitation and quantitative mass spectrometry analysis of recombinant U1-70K and deletions lacking the C-terminal LC domain(s). A network-driven approach resolved functional classes of U1-70K interacting proteins that showed dependency on the U1-70K LC domain(s) for their interaction. This included structurally similar RBPs, such as LUC7L3 and RBM25, which require their respective mixed charge domains for reciprocal interactions with U1-70K and for participation in nuclear RNA granules. Strikingly, a significant proportion of RBPs with mixed charge domains have elevated insolubility in AD brain proteome compared to controls. Furthermore, we show that the mixed charge LC domain of U1-70K can interact with Tau from AD brain. These findings highlight mechanisms for mixed charge domains in stabilizing RBP interactions and in potentially mediating co-aggregation with pathological Tau isoforms in AD.

## INTRODUCTION

The molecular processes that contribute to neurodegenerative diseases are not well understood. Recent observations suggest that numerous neurodegenerative diseases are promoted by the accumulation of RNA-binding protein (RBP) aggregates (1–3). This includes Alzheimer’s disease (AD), where pathological RNA-protein aggregates are often, but not exclusively, associated with Tau neurofibrillary tangles in brain (4). For example, U1 small nuclear ribonucleoprotein 70 kDa (U1-70K) and other core components of the spliceosome complex form detergent-insoluble aggregates in both sporadic and familial human cases of AD (5–7). Furthermore, RNA-seq analysis from AD and control brains revealed a significant accumulation of unspliced pre-mRNA disease related transcripts in AD consistent with a loss of U1-spliceosome function (7,8). Currently, our knowledge of the specific mechanisms underlying U1-70K aggregation is limited. This has proved to be a barrier to developing cellular models that would further our understanding of U1-70K and related RBP aggregation events in the pathogenesis of AD.

Supporting evidence indicates that a select group of RBPs are poised for aggregation because they self-assemble to form structures, including RNA granules (9), which are membrane-free organelles composed of RNA and RBPs (1,9–12). It has been proposed that RNA granules form via liquid–liquid phase separation (LLPS), which is driven by a dynamic network of multivalent interactions between structurally disordered low complexity (LC) domains (13–15) that have limited diversity in their amino acid composition (16). LLPS allows specific RBPs to concentrate and separate, leading to the formation of higher-order structures including oligomers, granules, and ultimately aggregates (16–18). Notably, several RBPs that aggregate in neurodegenerative disease contain LC domains, including TDP-43 and FUS (19–21). The LC domains found in TDP-43 and FUS mediate self-association, are necessary for RNA granule formation, and polymerize into amyloid-like aggregates (9,22,23). Mutations harbored within the LC domains of TDP-43 and FUS cause amyotrophic lateral sclerosis (ALS) and increased RNA granule stability, highlighting a critical role for LC domains in disease pathogenesis (24–26).

We recently reported that human AD brain homogenates induced the aggregation of soluble U1-70K from control brain and recombinant U1-70K, rendering it detergent-insoluble (5). The C-terminus of U1-70K, which harbors two LC domains (LC1 and LC2) was necessary for this aggregation (5). Furthermore, the LC1 domain (residues 231–308) of U1-70K was sufficient for robust aggregation, and through cross-linking studies was found to directly interact with insoluble U1-70K in AD brain homogenates (5). Collectively, these observations led to a hypothesis that pathological aggregation of RBPs in neurodegenerative diseases, including U1-70K, is driven by LC domains. However, unlike the prion-like Q/N-rich LC domains of TDP-43 and FUS, the LC1 domain of U1-70K contains highly repetitive complementary basic (R/K) and acidic (D/E) residues. These structurally unique motifs were originally described by Perutz (27), who proposed their ability to self-assemble and form higher-order structures termed polar zippers (27). Currently the physiological role of polar zipper motifs in U1-70K and other RNA binding proteins is unclear, and understanding their role in protein-protein interactions may shed light on the mechanisms underlying RBP aggregation and their association with Tau in AD (7,28).

Here we report that the mixed charge LC1 domain of U1-70K shares many of the same properties of the Q/N-rich LC domains found in TDP-43 and FUS, despite having a vastly different amino acid composition. These properties include the ability to self-assemble into high molecular weight oligomers and associate with nuclear granules in cells. To analyze the functional roles for the LC domains in U1-70K, we performed co-immunoprecipitation of recombinant U1-70K and serial deletions lacking one or both LC domains followed by quantitative proteomic analysis. Using a network-based bioinformatic approach we mapped classes of U1-70K interacting proteins that showed a dramatic reduction in their association with U1-70K in the absence of the LC1 domain. Remarkably, this revealed a group of functionally and structurally similar RBPs that also contained mixed charge domains analogous to the LC1 domain in U1-70K. These included LUC7L3 and RBM25, which we confirm also require their respective mixed charge domains for reciprocal interactions with U1-70K and for proper nuclear RNA granule association in cells. Furthermore, global analysis of the AD detergent-insoluble proteome revealed elevated levels of mixed charge RBPs within AD brain compared to controls. Finally, we show that the LC1 domain of U1-70K can interact with Tau from AD brain, which supports a hypothesis that mixed charge structural motifs on U1-70K and related RBPs could mediate cooperative interactions with Tau in AD.

## RESULTS

### The LC1 domain of U1-70K is necessary and sufficient for self-association in cells

The U1-70K LC1 domain (residues 231–308) is necessary and sufficient for robust aggregation in AD brain homogenates (5), yet it remains unknown if this domain is required for endogenous U1-70K self-association under physiological conditions. To test this hypothesis, we over-expressed full-length recombinant GST-fused and Myc-tagged U1-70K (rU1-70K) in HEK293 cells with serial deletions lacking one or both LC domains followed by co-immunoprecipitation (co-IP) and western blot analysis (**Fig. 1A**). Full-length rU1-70K (WT) and the ΔLC2 mutant co-immunoprecipitated endogenous U1-70K (~55 kDa), while rU1-70K mutants lacking the mixed charge LC1 domain (ΔLC1 and ΔLC1+2) displayed a dramatic impairment in their ability to co-IP native U1-70K. Together, these observations indicate that the LC1 domain is necessary for U1-70K self-association. To determine if the LC1 domain is sufficient for self-association, co-IPs were performed following the over-expression of alternative rU1-70K truncations, which included the N-terminus expressed alone (residues 1–99), the N-terminus and RNA recognition motif (residues 1–181), LC1, and the LC2 domain expressed in isolation (**Fig. 1B**). Only the LC1 domain was sufficient for appreciable self-association with native U1-70K. Furthermore, this interaction was likely not influenced by the presence of RNA, as treatment of the lysates with RNase prior to co-IP did not impair the ability of rU1-70K to interact with native U1-70K (**Fig. 1C**). Thus, our findings support that the LC1 domain is necessary and sufficient for U1-70K self-association, and that this interaction is predominantly RNA-independent.

**Figure 1.**
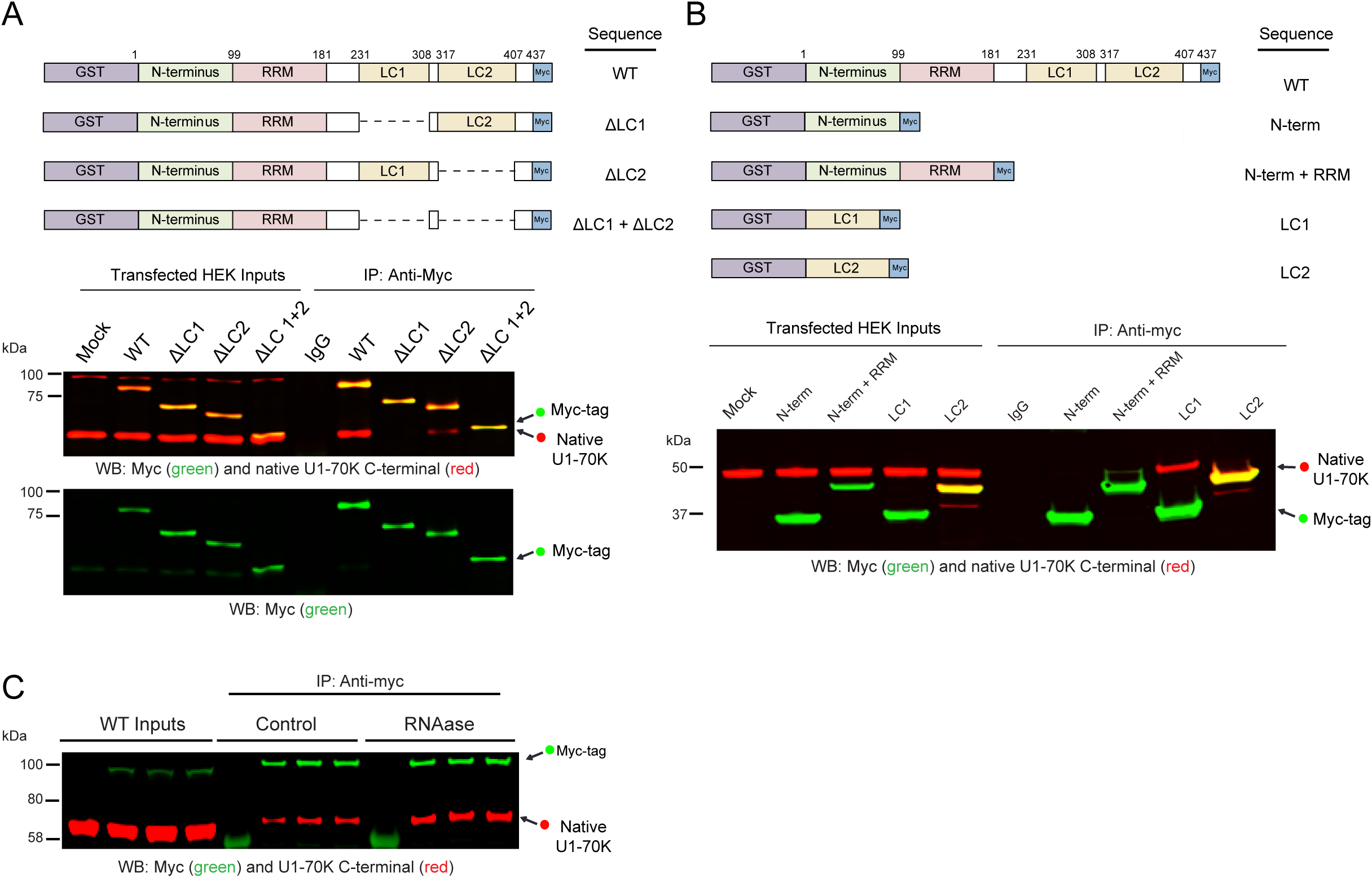
The LC1 domain of U1-70K is necessary and sufficient for self-association in cells. **A)** Full-length (WT) recombinant GST-fused and Myc-tagged U1-70K (rU1-70K) and serial deletions lacking one or both LC domains (ΔLC1 and ΔLC2, and ΔLC1+ΔLC2) were over-expressed in HEK293 cells and immunoprecipitated (IP) with anti-Myc antibodies. IP with a non-specific IgG was also performed from mock transfected cells as a negative control. Western blot for recombinant myc-tagged proteins (green) and native U1-70K (red) are shown for both the inputs and co-IPs (A, bottom panels). **B)** Full-length WT and rU1-70K truncations including the N-terminus (1–99 residues) alone, the N-terminus and RRM (1–181 residues), LC1 alone (231–308 residues) and the LC2 domain alone (317–407). IP with a non-specific IgG was also performed from mock transfected cells as a negative control. Western blot for recombinant myc-tagged proteins (green) and native U1-70K (red) are shown for both the inputs and co-IPs. **C)** WT rU1-70K was immunoprecipitated from untreated and RNAase (50 ng/uL) treated lysates followed by western blot for the myc-tag recombinant protein (green) and native U1-70K (red).

### The U1-70K LC1 domain oligomerizes in vitro

Although the LC1 domain of rU1-70K was deemed sufficient to interact with native Mixed Charge RNA binding proteins aggregate in AD U1-70K in cells, it was unclear whether this interaction is direct or facilitated by indirect interactions with additional RBPs. To determine if the LC1 of U1-70K can directly self-associate, we performed blue native gel polyacrylamide gel electrophoresis (BN-PAGE) of the GST-purified LC1 (residues 231–310) and N-terminal domain (residues 1–99) of rU1-70K; the latter was unable to interact with native U1-70K in cells (**Fig. 1B**). In contrast to SDS-PAGE, which resolves proteins under denaturing conditions, BN-PAGE is used to determine native protein complex masses, including high molecular weight oligomeric states and to identify physiological protein– protein interactions (29). Under the denaturing conditions of SDS-PAGE (**Fig. 2A**), both the LC1 and N-terminal domain have equivalent molecular weights (~65 kDa) compared to purified GST (~20 kDa). However, under native conditions (**Fig. 2B**), the LC1 domain formed dimers, trimers, tetramers and high-molecular weight oligomers that were in the megadalton MW range (>1,236 kDa). In contrast, the N-terminal domain mainly existed in the monomeric and dimeric state with some evidence of lower abundance high-molecular weight oligomers; GST alone was almost exclusively monomeric (**Fig. 2B**). These complexes became more readily visible following the transfer to a membrane and western blot analysis with Myc antibodies (**Fig. 2C**). Remarkably, a higher proportion of the LC1 domain formed dimers (*n*=2) and tetramers (*n*=4) compared to trimers (*n*=3), suggesting that dimer intermediates are favored over trimer intermediates for tetramer formation (**Fig. 2D**). These *in vitro* findings demonstrate that the LC1 domain can directly self-associate to form oligomers, including high molecular weight species, which implicates direct LC1-LC1 interactions as a mechanism of U1-70K self-association.

**Figure 2.**
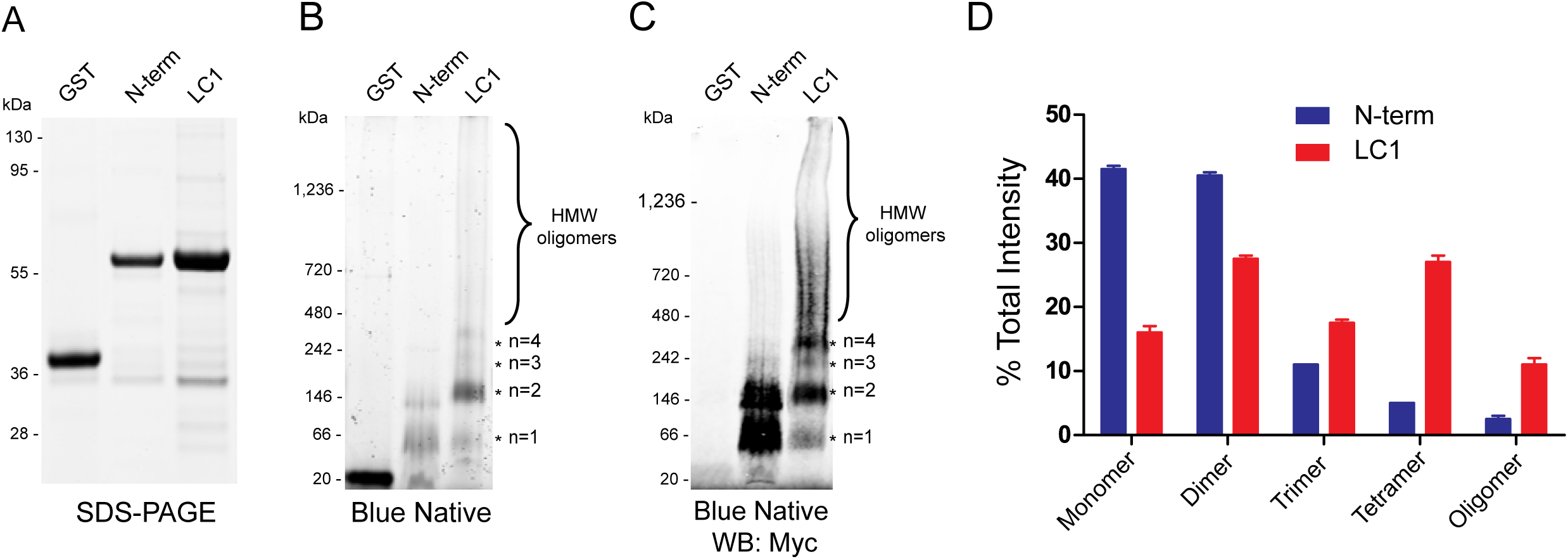
The LC1 domain of U1-70K directly self-interacts and oligomerizes *in vitro*. **A)** SDS-PAGE of GST alone, GST purified N-terminal domain and GST purified LC1 domain of rU1-70K. Both the LC1 and N-terminal domain have equivalent molecular weights (~65 kDa) while GST alone is (~20 kDa). **B)** Blue native gel polyacrylamide gel electrophoresis (BN-PAGE) of GST alone, the N-terminal domain and the LC1 domain of rU1-70K, respectively. The LC1 domain formed higher molecular weight species (*) consistent with dimers (~130 kDa), trimers (~195 kDa), tetramers (~260 kDa) and high-molecular weight (HMW) oligomers (>400 kDa). **C)** Western blot detection of blue native complexes using Myc antibodies. **D)** Densitometry of monomeric, dimeric, trimeric and HMW species of the N-terminal domain (blue) and LC1 domain (red) of rU1-70K. Each form is represented as the fraction of total signal in each sample analyzed in technical replicate (*n*=2). Error bars represent the standard deviation (s.d.).

### The U1-70K LC1 domain is necessary and sufficient for robust nuclear granule localization

Our data highlights that a key role for the U1-70K LC1 domain is self-association and oligomerization. However, it is not well established whether the LC1 domain influences nuclear localization and granule formation in cells. To address this question, full-length rU1-70K or mutants lacking one or both LC domains (**Fig. 1A**) were over-expressed in HEK293 cells followed by subcellular biochemical fractionation into nuclear and cytoplasmic pools (**Fig. 3A-B**). The rU1-70K mutants containing a LC1 domain (WT and ΔLC2) partitioned mainly to the nuclear fraction (~75% nuclear), but mutants lacking the LC1 domain (ΔLC1 and ΔLC1+2) were equally distributed between nucleus and cytoplasm, indicating a significant impairment of nuclear localization. These biochemical findings were further supported by immunocytochemistry, which showed that rU1-70K mutants lacking the LC1 domain displayed diffuse expression patterns in both the nucleus and cytoplasm compared to WT and ΔLC2 proteins. (**Fig. 3C**). Given that the LC1 domain in isolation can directly self-associate and oligomerize (**Fig. 2**), we sought to determine if the LC1 domain was necessary for RNA granule localization in cells (**Fig. 3C**). As expected, full-length rU1-70K protein localized to nuclear granules, in agreement with previous studies (30–32). Although the nuclear granule localization of mutants lacking the LC1 domain was diminished, the ΔLC2 mutant retained nuclear granule localization, supporting the necessity of the LC1 domain for subnuclear granule localization. Furthermore, the LC1 domain alone was found to colocalize with native U1-70K in nuclear granules (**Fig. 3D**), consistent with the ability of the LC1 domain to interact with native U1-70K from cell lysates (**Fig. 1D**). Collectively, these results demonstrate that the LC1 domain is important for U1-70K subcellular nuclear localization, and suggests a role for LC1 mediated intermolecular interactions as a mechanism of nuclear RNA granule formation.

**Figure 3.**
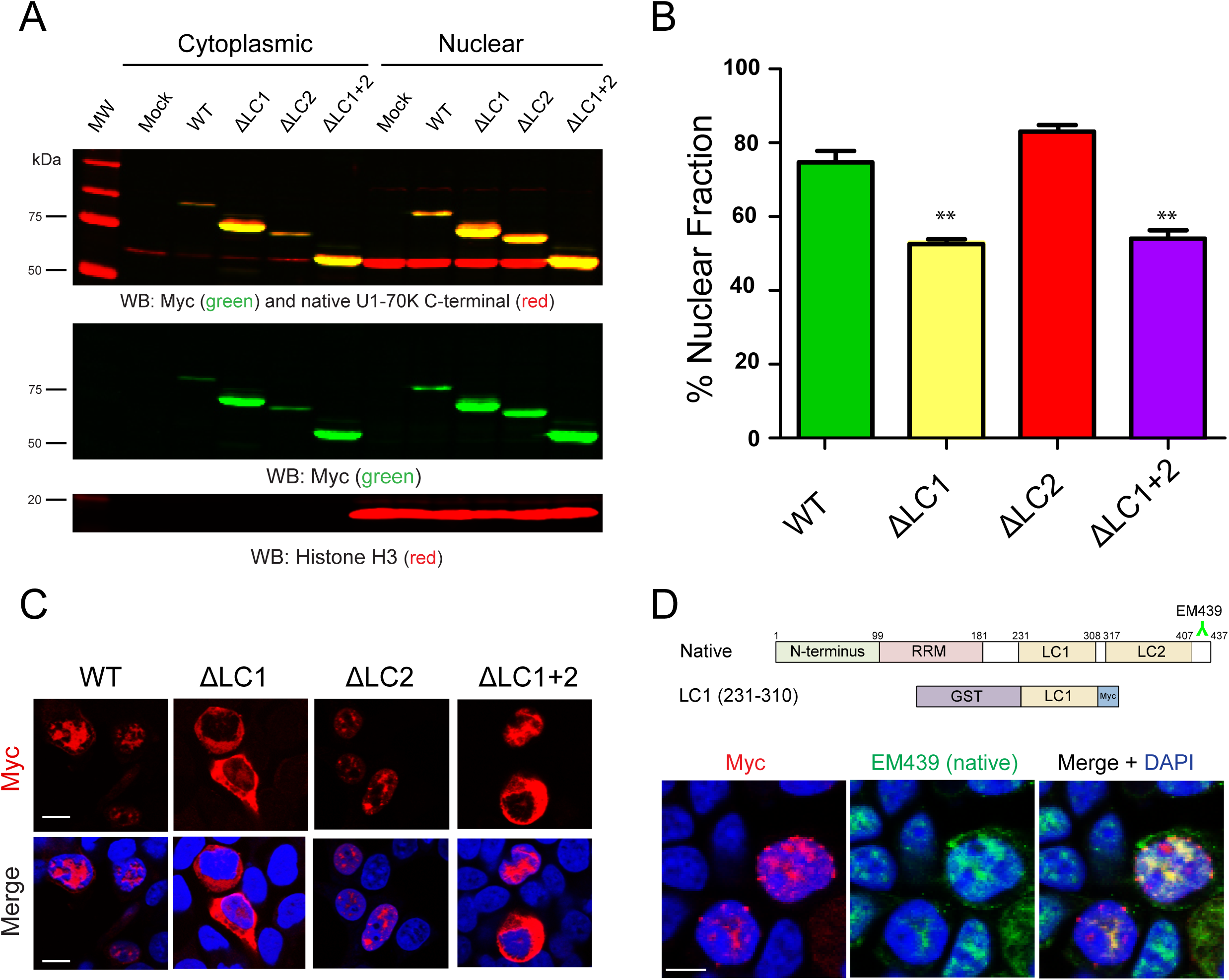
The LC1 domain is necessary and sufficient for robust nuclear granule formation. **A)** WT full-length rU1-70K or mutants lacking one or both LC domains were over-expressed in HEK293 cells. The cells were then fractionated into nuclear and cytoplasmic pools followed by western blot analysis for both recombinant myc-tagged proteins (green) and native U1-70K (red). Western blots for histone H3 (bottom panel) was used as a positive control in the nuclear fraction. **B)** Densitometry analysis was performed to calculate the levels of cytoplasmic and nuclear rU1-70K and mutants and % nuclear intensity for each rU1-70K protein is reported. Each experiment was performed in biological triplicate (*n*=3) with error bars representing the standard deviation (s.d.). Both the ΔLC1 and ΔLC1+2 rU1-70K fragments were significantly less nuclear than full length rU1-70K (** p-value<0.01). **C)** Immunocytochemistry for WT rU1-70K and mutants that lacked the LC1, LC2 or both LC domains was performed and visualized by confocal microscopy. Scale bar equates to 10 μM. **D)** Over-expression of the rU1-70K LC1 domain alone (red) resulted in robust nuclear granule formation and sequestration of native U1-70K (green). The EM439 detects and extreme C-terminal epitope not present in the LC1 rU1-70K fragment, which allows discrimination between the recombination protein and the native U1-70K. DAPI stained nuclei are shown in blue. Scale bar equates to 10 μM.

### Protein-Protein interaction network analysis resolves functionally distinct classes of U1-70K interacting proteins

To further assess the physiological function of the LC domains of U1-70K, we performed co-immunoprecipitations of full-length rU1-70K and various rU1-70K mutants lacking either or both LC domains from HEK293 cells followed by liquid chromatography coupled to tandem mass spectrometry (LC-MS/MS) to identify co-purified interacting proteins. Each co-IP was performed in biological quadruplicate (*n*=4) and an equal number of mock IPs were performed using a non-specific immunoglobulin (IgG) as a negative control. Protein abundance was determined by peptide ion-intensity measurements across LC-MS/MS runs using the label-free quantification (LFQ) algorithm in MaxQuant (33). In total, 45,223 peptides mapping to 3,458 protein groups were identified. However, one limitation of data-dependent LFQ proteomics methods is the inherent missing data (i.e. missing protein identifications or abundance values), especially for low abundance proteins (34). Thus, proteins with ten or more missing values across the 20 samples were not included in the subsequent bioinformatic analysis. To limit the number of non-specific interactors, proteins with less than a 1.5-fold enrichment over IgG were not considered. This resulted in the final quantification of high-confidence interactors falling into 716 protein groups mapping to 713 unique gene symbols (**Supplemental Table 1**).

WeiGhted Co-expression Network Analysis (WGCNA) is typically used for large-scale transcriptome and proteome datasets to categorize gene products into biologically meaningful complexes, molecular functions and cellular pathways (35). Here, we sought to leverage co-enrichment patterns to better classify protein-protein interactions (PPIs) across WT and rU1-70K deletions to assess if specific classes of proteins selectively favor interactions with the LC domains. In WGCNA, correlation coefficients between each protein pair in the dataset is calculated and groups of highly correlated proteins are segregated into modules (36). In our dataset, a total of 7 modules were defined (**Supplemental Fig. 1** and **Fig. 4A**). These modules range from 292 proteins in turquoise to 19 in black (**Supplemental Table 1**). The premise of co-expression or in this case protein co-enrichment analysis is that the strong correlation between two or more proteins is indicative of a physical interaction, functional relationship, and/or co-regulation. We therefore hypothesized that following a co-IP for rU1-70K, specific modules would reflect biologically relevant PPIs, and thus highlight distinct complexes. As expected, modules were significantly enriched for biologically meaningful gene ontologies (GO) as well as established cellular functions and/or organelles as determined by GO-Elite (**Table 1**).

**Figure 4.**
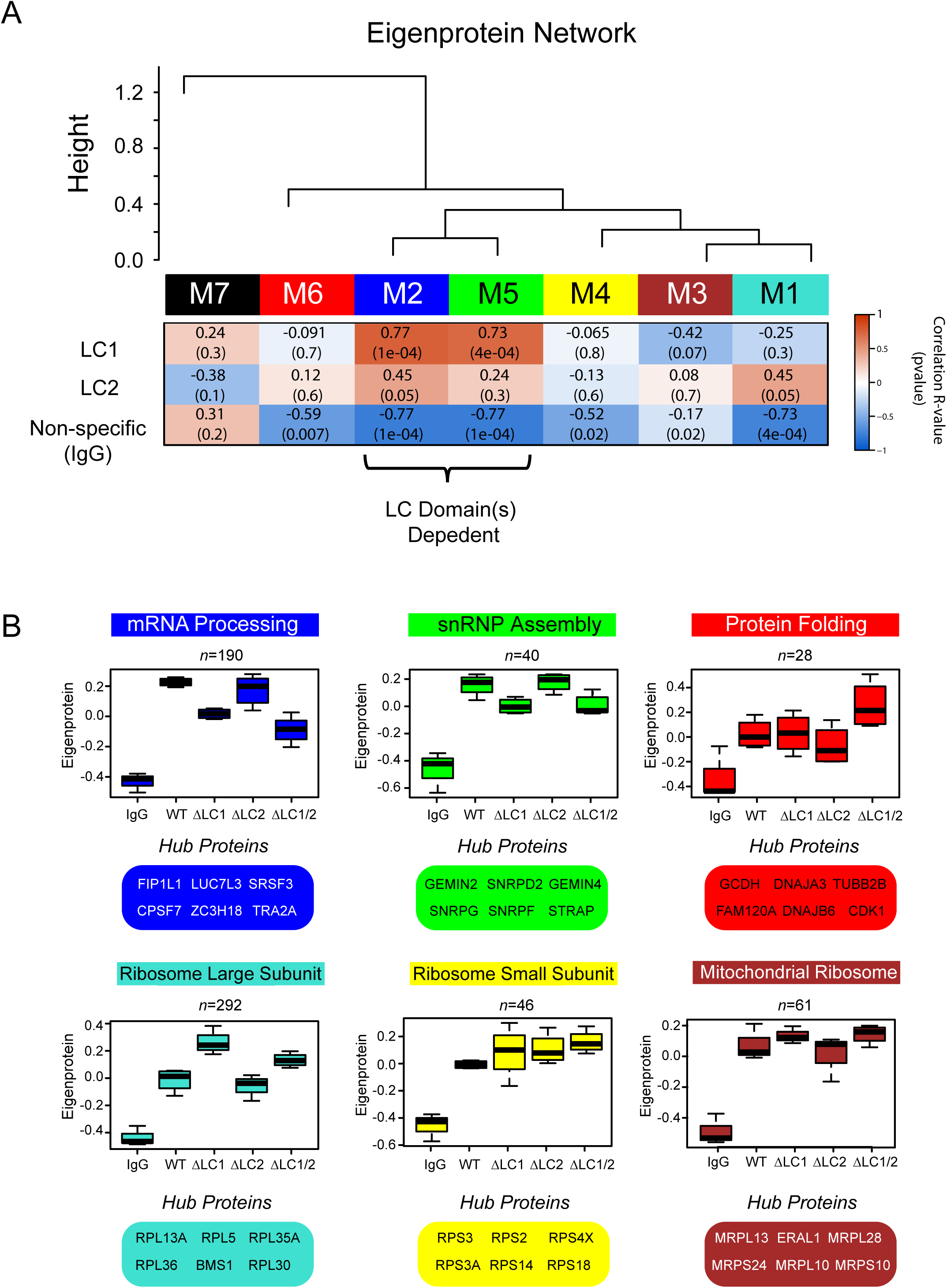
Correlation network analysis resolves distinct modules of U1-70K interacting proteins that differ in their association with the LC1 domain. **A)** WGCNA organized proteins defined by dendrogram branch cutting (**Supplemental Figure 1**) measured across all co-IP samples (*n*=716) into modules (M1-M7) that represent clusters of proteins defined by their correlation to each other across the five co-IP conditions analyzed (IgG, WT rU1-70K and deletions ΔLC1, ΔLC2, and ΔLC1+ΔLC2). Listed in the heatmap are bicor correlations and p-values defining relationship between module eigenprotein level and rU1-70K protein (defined as 0-IgG, 1-ΔLC1, and 3-ΔLC2). **B)** Eigenproteins, which correspond to the first principal component of a given module and serve as a summary expression profile for all proteins within a module, are shown for 6 modules generated by WGCNA. Box plots with error bars beyond the 25th and 75th percentiles are shown for all four groups (IgG, WT, ΔLC1, ΔLC2, and ΔLC1+2). Hub proteins for each of these modules are also highlighted below for each module.

**Table 1.**
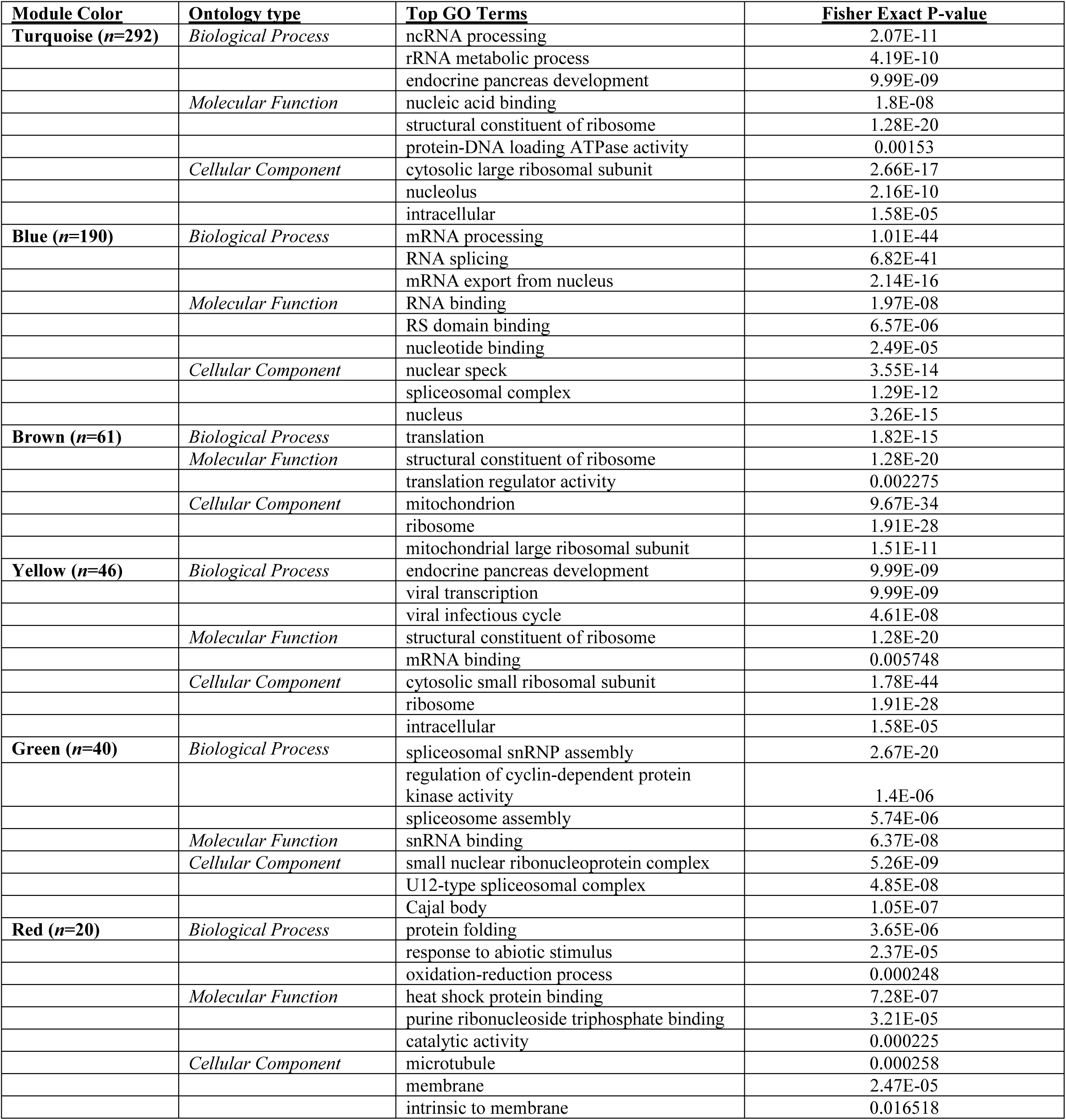
U1-70K Protein-Protein Interaction Network Generates Modules Enriched with Specific Gene Ontology (GO) terms

Each module has an abundance profile for all member proteins across the rU1-70K co-IP conditions, termed the eigenprotein (**Fig. 4A-B**). Notably, six (M1-M6) of the seven modules showed a significantly higher level of co-enrichment in the WT and rU1-70K deletions compared to the IgG negative controls, indicative of specific interactions for members of these modules (**Fig 4A-B**). In contrast, M7 (black) had essentially equivalent levels across all conditions and was the only module with positive correlation to the non-specific IgG. Therefore, it was considered non-specific and not used for further analysis **(Fig. 4A)**. The protein interactors with reduced affinity for rU1-70K following LC1 deletion (e.g. ΔLC1 and ΔLC 1/2), included members of both the mRNA processing (blue) and snRNP assembly (green) modules (**Table 1**). Therefore, the LC1 dependent interactors are defined by membership to these modules. In contrast, modules enriched with large ribosomal subunit components (turquoise) and mitochondrial ribosome subunits (brown) displayed increased levels following deletion of one or both LC domains, suggesting that the LC1 domain negatively regulates their interactions with U1-70K (**Table 1 and Fig. 4A-B**). Finally, the modules enriched with proteins involved in protein folding (red) and the small ribosomal subunit (yellow) showed little difference in expression across the co-IP conditions, suggesting that these protein interactions are mainly with the N-terminus and/or RRM domain of U1-70K. The kME score quantifies how well a given protein pattern matches that of the module eigenprotein, with high scores approaching one signifying a high correlation (37). The hub proteins, with the highest correlation to the module abundance profile (i.e., eigenprotein), are highlighted in (**Fig. 4B)**. These findings demonstrate that a weighted PPI network analysis of the U1-70K interactome successfully resolved biologically and structurally distinct complexes.

### Confirmation of U1-70K LC1 dependent interacting proteins

To validate and extend the module assignments, we performed both *in silico* and biochemical analysis. First, to visualize the relationships among modules with an independent clustering method, the T-Distributed Stochastic Neighbor Embedding (tSNE) algorithm was used to map the relatedness of proteins of top module members. The tSNE analysis largely agreed with and confirmed the module assignment, whereby the majority of proteins clustered with their own module members as assigned by WGCNA (**Fig. 5A**). The tSNE analysis also allows for visualization of module relatedness, with distance between proteins and clusters representing similarity of co-enrichment, with similar modules in close proximity to each other and dissimilar modules further apart. For example, modules involved in translation cluster together (brown and turquoise) while those involved in mRNA processing (blue) and snRNP assembly (green) formed a separate cluster.

**Figure 5.**
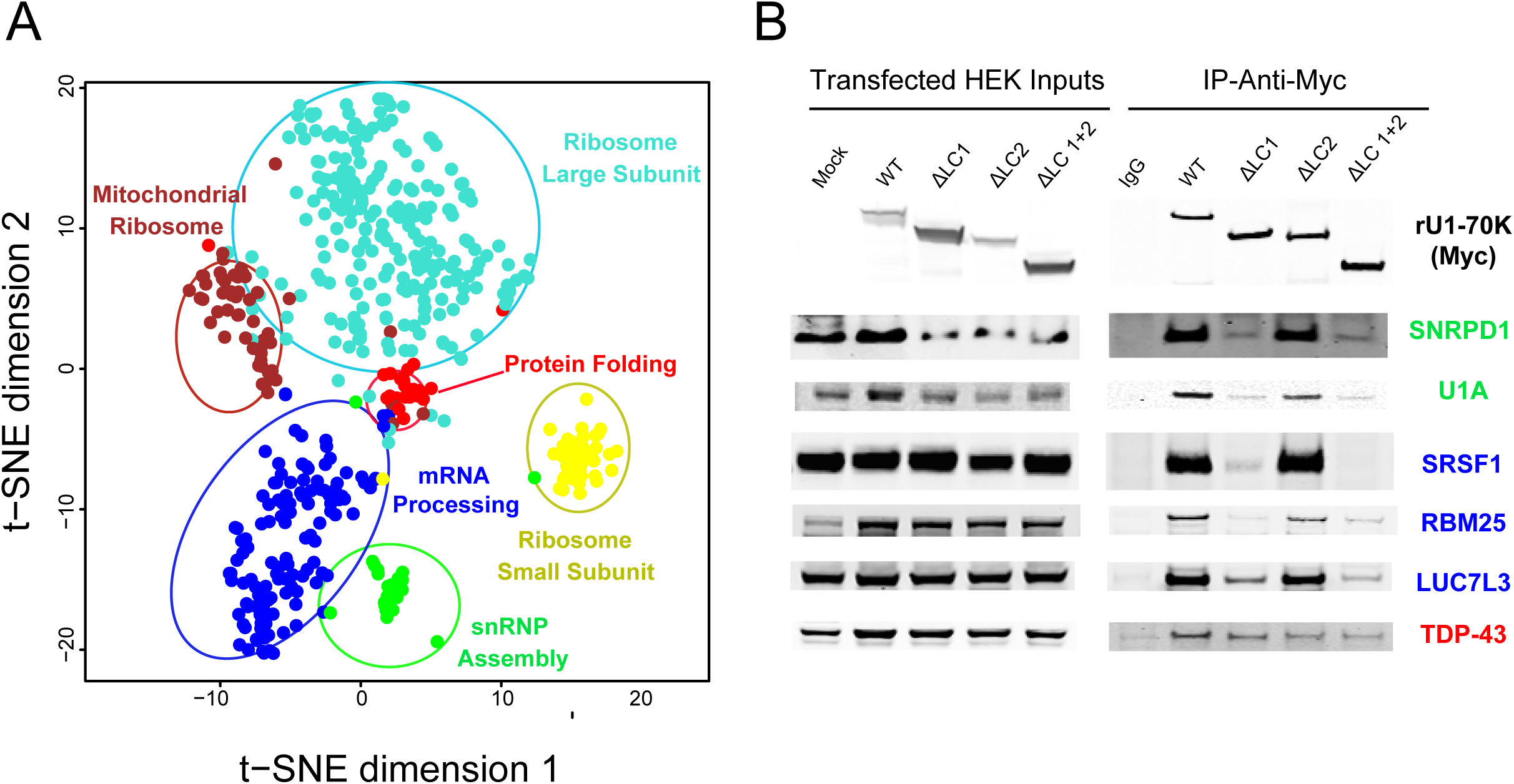
Confirmation of U1-70K interacting partners that favor interactions via the LC1 domain. **A)** To visualize the relationship between modules and validate the WGNCA results, the T-Distributed Stochastic Neighbor Embedding (tSNE) algorithm was used to map the relatedness of proteins with a kME score of 0.5 or greater, which is a measure of intramodular connectivity (kME), defined as the Pearson correlation between the expression pattern of a protein and the module eigenprotein. The tSNE analysis overlaid with module assignments determined by WGCNA allows for visualization of module relatedness, with distance between proteins representing similarity of co-enrichment, with the more similar clusters of proteins (related modules) in closer proximity to each other compared to dissimilar modules. Proteins with similar co-enrichment across the rU1-70K co-IPs are highly correlated to one another and are related to modules with distinct biological functions (**Table 1**). **B)** Western blot analysis to validate the proteomic results and module assignment were performed for select interactors of the snRNP assembly (SNRPD1 and U1A), mRNA processing (SRSF1, RBM25 and LUC7L3), and Protein folding (TDP-43) modules following co-IP across the five experimental conditions (IgG, WT rU1-70K and deletions ΔLC1, ΔLC2, and ΔLC1+ΔLC2).

To experimentally validate the module assignments biochemically, western blots were performed for select rU1-70K interactors of the snRNP assembly module (SNRPD1 and U1A), mRNA processing module (SRSF1, RBM25 and LUC7L3) and the protein folding module (TDP-43) (**Fig. 5B**). Members of both the mRNA processing and snRNP assembly modules had higher abundance in the WT and ΔLC2 co-IPs compared to that of the IgG, ΔLC1, and ΔLC1+2 IP samples. This mirrored the pattern observed for the blue and green eigenprotein values confirming the proteomic findings (**Fig. 4B**). In contrast, TDP-43, showed a similar level of interaction across WT rU1-70K and the various deletions, consistent with TDP-43 being an N-terminal interactor of U1-70K and not influenced by the absence of the disordered LC domains.

### The mRNA processing module is enriched with structurally similar RNA-binding proteins harboring mixed charge domains

U1-70K interacting proteins mapping to the mRNA processing and snRNP assembly modules are related by their affinity for the LC1 domain, yet they contain RNA binding proteins with distinct biological functions (**Table 1**). For example, all U1 snRNP components and assembly factors including U1A, U1C, Sm proteins, and the SMN complex are enriched in the green module. The SMN complex is responsible for loading the Sm proteins onto the snRNA scaffold, a critical step in U1 snRNP assembly (38). This module also contains the components of the 7SK snRNP which regulates snRNA transcription and is present in Cajal bodies, a site of U1 snRNP maturation (39,40). In contrast, the mRNA processing module (blue) is enriched with proteins associated with RNA splicing, polyadenylation, mRNA export and "nuclear specks", the latter being an analogous term for splicing speckles (41). However, it is their respective association with rU1-70K mutants that discriminates the mRNA processing and snRNP modules members. For example, proteins involved in snRNP assembly are less influenced by the loss of the LC2 domain, yet association of members of the mRNA processing module are affected, suggesting that proteins involved in granule/speckle assembly interact in part via both LC domains of U1-70K, whereas core spliceosome assembly factors do not favor interactions with the LC2 domain (42,43).

Our observations also revealed that several members of the mRNA processing module (blue) contain stretches of highly repetitive complementary basic (R/K) and acidic (D/E) residues, analogous to the LC1 domain of U1-70K (**Fig. 6A**), that we refer to as mixed charge domains. To examine the relationship between this sequence similarity and U1-70K interacting proteins, a list of LC1-like (i.e., mixed charge) proteins was created using the Uniprot protein Blast feature. Many proteins (n=255) in the proteome were determined to have significant sequence overlap to the mixed charge LC1 domain of U1-70K. These included other members of the mRNA processing module such as RBM25, ZC3H13, DDX46 and LUC7L3 among others. Although not identical in length, sequence alignment highlights the similar stretches of highly repetitive complementary basic (R/K) and acidic (D/E) residues across these distinct gene products (**Fig. 6A**). Indeed, a one-tailed Fisher’s exact test revealed that the mRNA processing module was significantly enriched with proteins harboring LC1-like mixed charge domains generated from the Blast analysis (**Fig. 6B**). In contrast, a similar analysis comparing disordered RNA binding proteins harboring prion-like (Q/N-rich) domains (44), including TDP-43 and FUS, showed no enrichment in any of the modules of U1-70K interacting proteins. Notably, the members of the mRNA processing module also had a significant over-representation of nuclear proteins that selectively precipitate after treatment with biotinylated isoxazole (b-isox) (45). Many of these proteins have been shown to participate in RNA granule assembly and to form hydrogels *in vitro* (9,46). Collectively these results suggest that structurally similar mixed charge domains, analogous to the U1-70K LC1 domain, engage in protein-protein interactions, which are essential for nuclear granule assembly.

**Figure 6.**
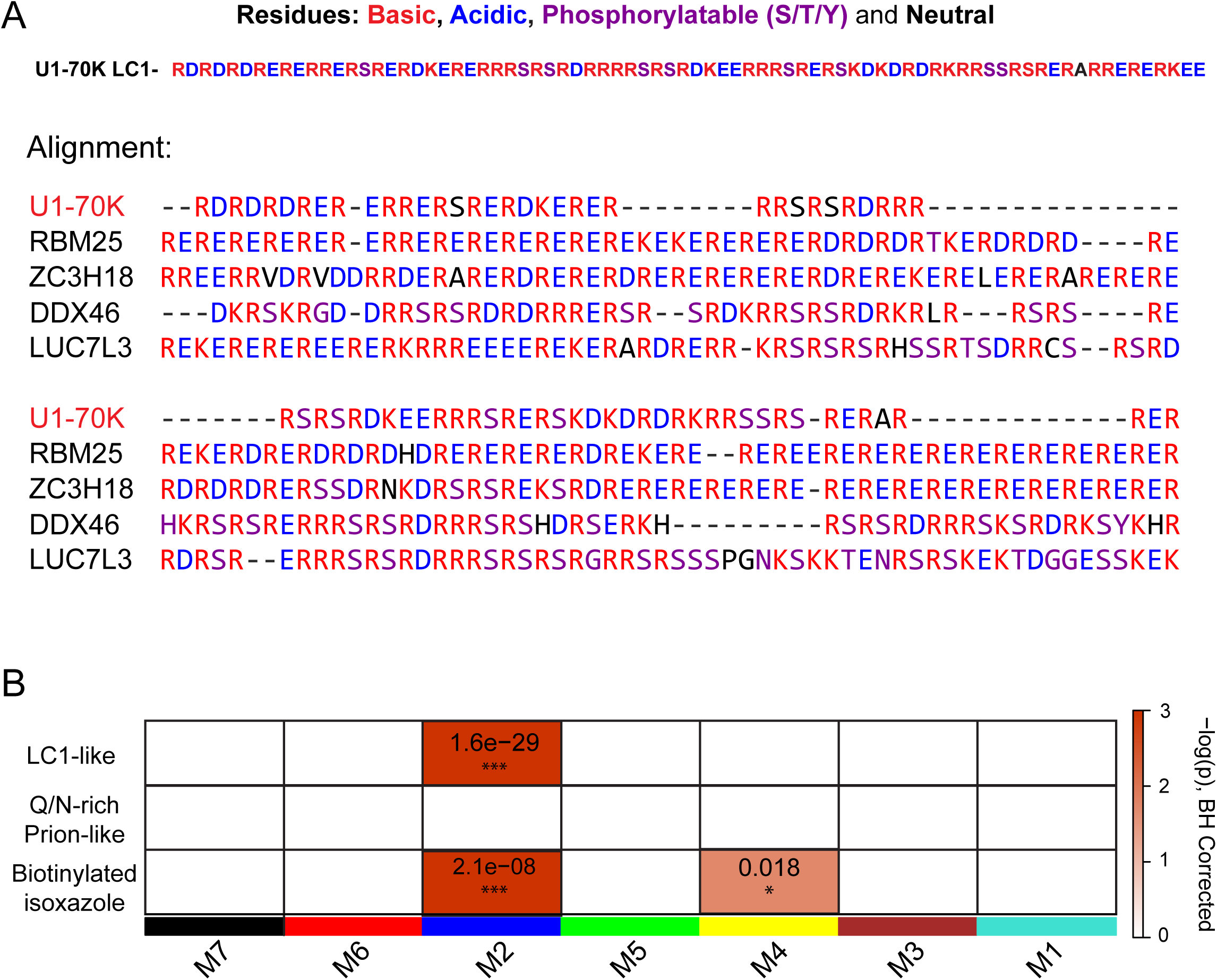
The mRNA processing module is enriched with structurally similar RNA-binding proteins harboring U1-70K LC1-like domains. **A)** The LC1 domain of U1-70K (residues 231–308) contains highly repetitive complementary basic (R/K) and acidic (D/E) residues. A list of 255 proteins that shared greater than 20% similarity to the LC1 domain of U1-70K (E-values less than 0.005) was created using the Uniprot protein Blast feature. Using clustal omega, an alignment was performed on the U1-70K LC1 domain and the four most structurally similar proteins to highlight the mixed charge nature of their sequence. **B)** A one-tailed Fisher’s exact test (FET) was used to assess structural overlap of LC1-like mixed charge proteins from Blast analysis with module membership for U1-70K interacting partners (upper panel). FET analysis was also performed using prion-like RNA binding proteins (middle panel) or proteins that were precipitated from nuclear extracts using biotin-isoxazole compound (bottom panel). Benjamin Hochberg corrected p-values (to control FDR for multiple comparisons) for the module enrichment is highlighted. Significance is demonstrated by the color scales, which go from 0 (white) to 3 (red), representing −log(p).

### Mixed charge domains in LUC7L3 and RBM25 are necessary for reciprocal interactions with U1-70K and nuclear RNA granule assembly

Based on their related structural and functional roles in RNA speckle assembly and affinity for the LC domains of U1-70K we asked whether members of the mRNA processing module co-localize with U1-70K in cells. Both RBM25 and LUC7L3 have mixed charge domains analogous to the LC1 domain of U1-70K with similarity E-values of 5.5E-27 and 1.8E-26, respectively, and amino acid overlap of 55.1% and 44.2%, respectively (**Fig 6A**). As expected, all three proteins were observed in nuclear granules (30,47,48), where U1-70K showed strong co-localization with LUC7L3 and RBM25 (**Fig. 7A**). Our current findings support that the LC1 domain is necessary and sufficient for U1-70K self-association and nuclear granule assembly. By extension we hypothesized that the mixed charge domains in LUC7L3 and RBM25 would similarly be important in mediating interactions with U1-70K and other structurally similar proteins. To test this possibility, we over-expressed full-length recombinant GST-fused and Myc-tagged rRBM25 or rLUC7L3 in HEK293 cells and their respective deletions lacking the mixed charge domains (ΔMC) and the mixed charge (MC) domains alone followed by IP and western blot analysis (**Fig. 7B**). The full-length rLUC7L3 and the MC domain were each able to co-IP endogenous LUC7L3, mirroring the self-association observed for U1-70K (**Fig. 1A**). Unfortunately, the rLUC7L3-ΔMC protein migrated at a similar molecular weight to endogenous LUC7L3, thus, we were unable to determine if the MC domain was necessary for self-association in cells. Full-length rLUC7L3 and the MC domain could also interact with endogenous U1-70K and RBM25 (**Fig. 7B**), whereas the rLUC7L3-ΔMC mutant was unable to support these interactions **(Fig. 7B)**. Similarly, full-length rRBM25 could interact with both endogenous U1-70K and LUC7L3, while the rRBM25-ΔMC mutant was unable to support these interactions (**Fig. 7C**). In contrast, the MC domain of rRBM25 was not sufficient to interact with U1-70K or LUC7L3, perhaps due to misprocessing, post-translation modifications, or size.

**Figure 7.**
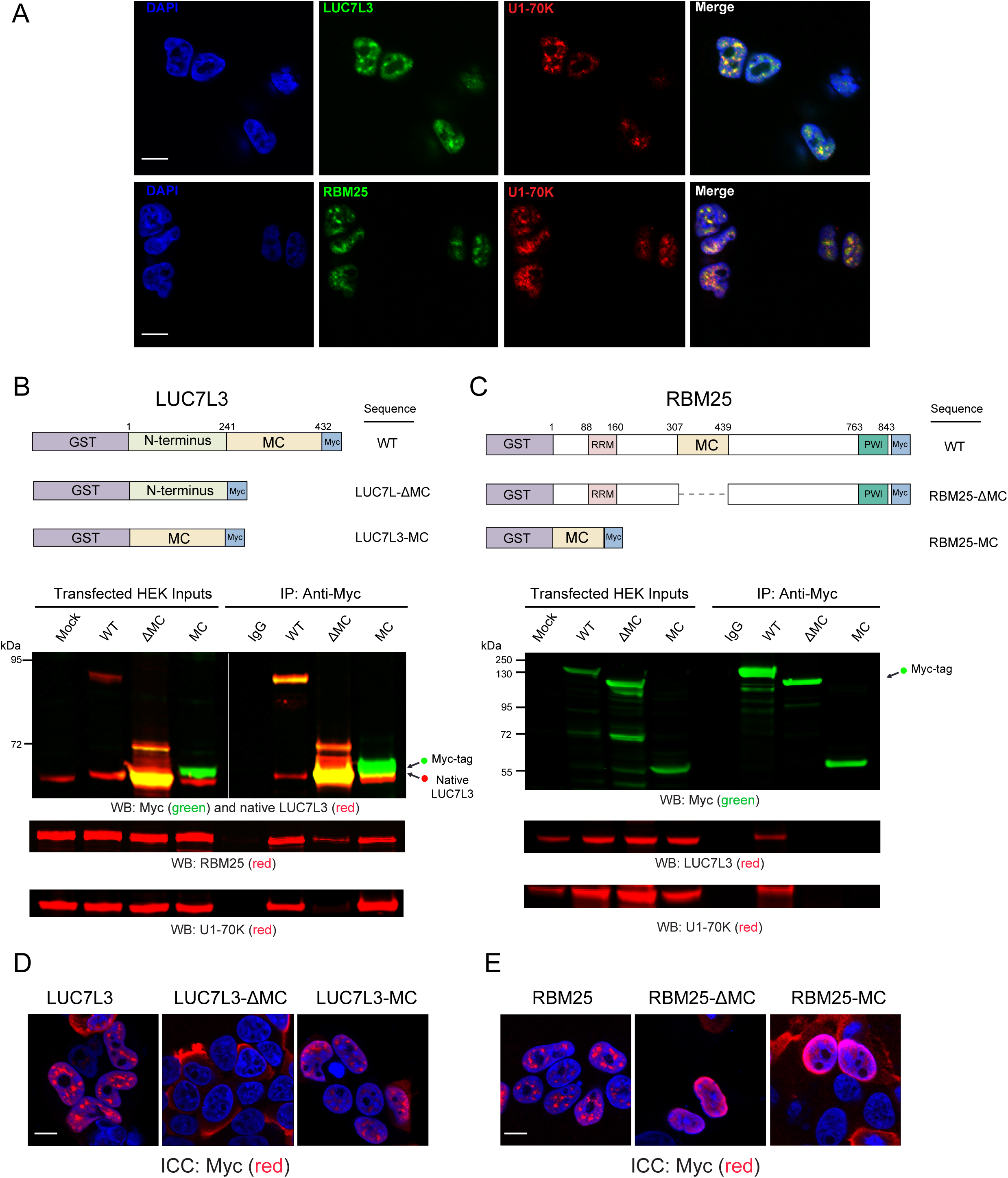
The mixed charge domains in LUC7L3 and RBM25 are necessary for reciprocal interactions with U1-70K and nuclear RNA granule assembly. **A)** Immunocytochemistry (ICC), to assess the co-localization of native U1-70K (green) with RBM25 (red) or LUC7L3 (red). DAPI stained nuclei are shown in blue. Scale bar equates to 10 μM. **B)** Full-length (WT) recombinant GST-fused and Myc-tagged LUC7L3 (rLUC7L3) and mutants lacking the mixed charge (MC) domain or the MC domain alone (upper panel) were over-expressed in HEK293 cells and immunoprecipitated (IP) with anti-Myc antibodies. IP with a non-specific IgG was also performed from mock transfected cells as a negative control. Western blot for recombinant myc-tagged proteins (green) and native LUC7L3 (red) are shown for both the inputs and co-IPs (bottom panels). Membranes were also re-probed for native U1-70K (red) or RBM25 (red). **C)** Full-length (WT) recombinant GST-fused and Myc-tagged RBM25 (rRBM25) and mutants lacking the MC domain or the MC domain alone (upper panel) were over-expressed in HEK293 cells and IP with anti-Myc antibodies. IP with a non-specific IgG was also performed from mock transfected cells as a negative control. Western blot for recombinant myc-tagged proteins (green) and native U1-70K (red) or LUC7L3 (red) are shown for both the inputs and co-IPs (bottom panels). **D)** Immunocytochemistry for full-length rLUC7L3, a mutant lacking the MC domain (LCU7L3-ΔMC) and the MC domain alone (LUC7L3-MC) were expressed in HEK293 cells and visualized by confocal microscopy. **E)** Immunocytochemistry for rRBM25, a mutant lacking the MC domain (RBM25-ΔMC) and the MC domain alone (RBM25-MC) were expressed in HEK293 cells and visualized by confocal microscopy. DAPI was used to visualize nuclei (blue), scale bars = 10µM for both panels D and E.

Given the role of the U1-70K LC1 domain in RNA granule formation, we sought to determine if the MC domains of LUC7L3 and RBM25 influenced nuclear localization to granules (**Fig. 7D and E**). Both full-length rLUC7L3 and rRBM25 localized to nuclear granules by immunocytochemistry, consistent with their endogenous LUC7L3 and RBM25 expression pattern in cells (**Fig. 7**). However, the rLUC7L3-ΔMC was observed diffusely localized to the cytoplasm, whereas rRBM25-ΔMC remained mainly in the nucleus, but did not localize to nuclear granules. Consistent with U1-70K, the MC domain of rLUC7L3 was sufficient to localize to nuclear granules, likely due to its interactions with U1-70K and other mixed charge RBPs. However, the MC domain of rRBM25, which was unable to interact by co-IP with mixed charge RBPs, did not form nuclear granules in the cells. Taken together, our findings suggest a shared functional role for mixed-charge domains in stabilizing protein-protein interactions, that likely play a role in nuclear granule assembly.

### RNA binding proteins with mixed charge domains have enhanced insolubility in AD brain

Based on the ability of U1-70K to aggregate in AD brain homogenate, and the key role of the LC1 domain in U1-70K oligomerization *in vitro*, we hypothesized that proteins harboring similar MC domains would preferentially aggregate in AD brain. To test this hypothesis, we assessed the distribution of insoluble proteins with MC domains in a recently published comprehensive analysis of the sarkosyl-insoluble proteome (*n*=4,643 proteins quantified) from individual control and AD cases (49). Protein ratios for all pairwise comparisons (i.e., control vs. AD) were converted into log_2_ values, and the resulting histogram fit to a normal Gaussian distribution (**Fig. 8A**). Compared to the normal distribution of all proteins in the AD insoluble proteome (purple histogram), quantified mixed charge proteins (yellow histogram) showed a global shift towards insolubility in AD (**Fig. 8A**). This increase was found to be significant using one-tailed Fisher exact test (p-value=2.028868e-09). Consistently, mixed charge proteins that fell within the top 10^th^ percentile (*n*=28) were significantly elevated in AD cases compared to controls, similar to Aβ and tau levels (**Fig. 8B and C**). Strikingly, 68% of the AD enriched mixed charge proteins, including LUC7L3, within the top 10^th^ percentile were members of the blue module from rU1-70K interactome studies **(Fig. 4**), with shared functions in RNA binding, splicing and processing (**Fig. 8D and E**). Collectively, these data indicate that RBPs with MC domains have a higher likelihood of insolubility and aggregation in AD brain.

**Figure 8.**
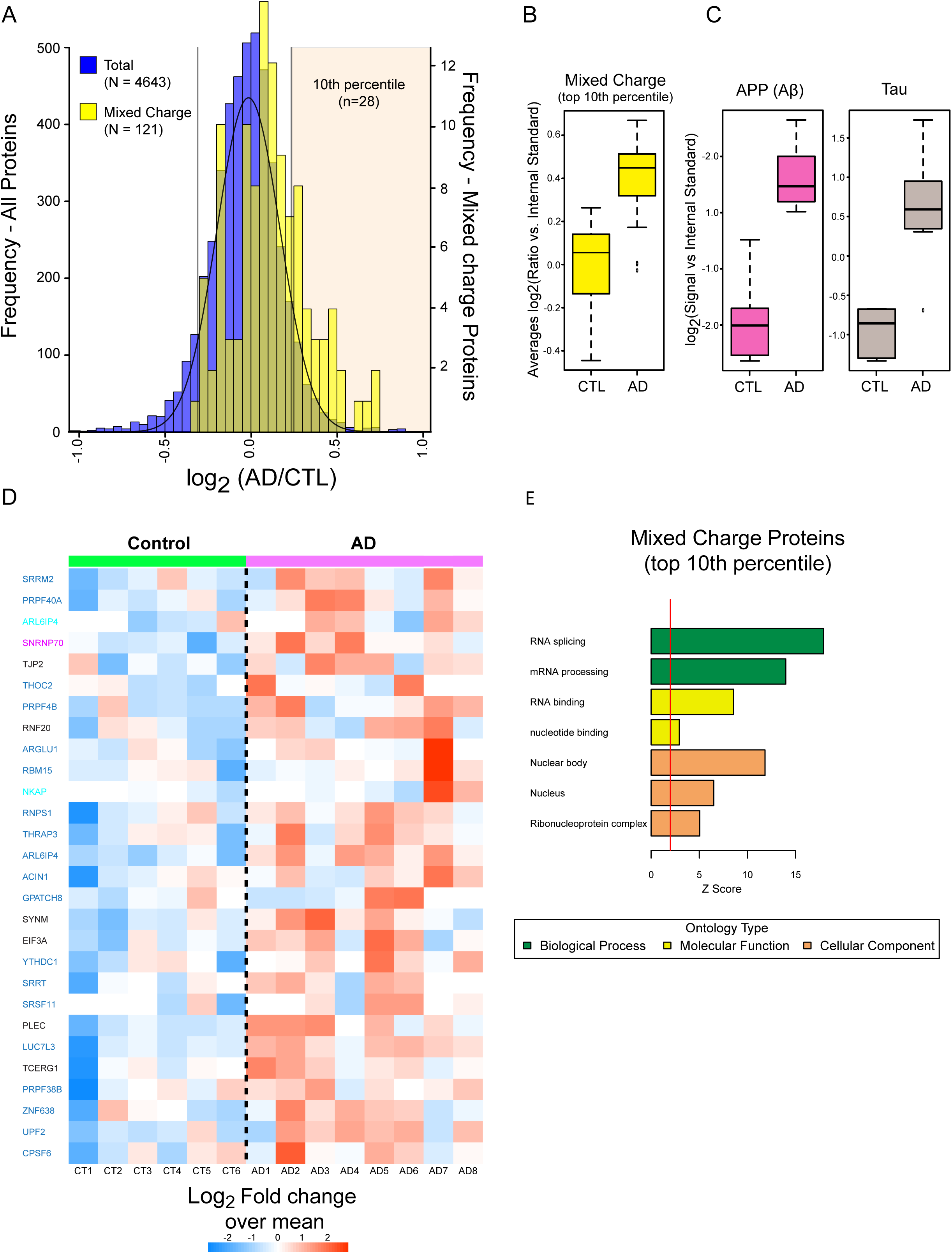
RNA binding proteins with mixed charge domains have increased insolubility in AD brain. **A)** Histogram of average log_2_ ratios (AD/control) from the control (*n*=6) and AD (*n*=8) brain detergent insoluble fractions. Protein ratios for all pairwise comparisons (i.e., control vs. AD) were converted into log_2_ values, and the resulting histogram fit to a normal Gaussian distribution. Compared to the normal distribution of all proteins in the AD insoluble proteome (purple histogram), quantified mixed charge proteins (*n* =112 yellow histogram) showed a global shift towards insolubility in AD. The 28 proteins with MC domains that fell above the ninetieth percentile (top 10% percentile) of the total population distribution are significantly overrepresented (Fisher exact p-value 2.0e-09). **B)** Box plot of the mixed charge proteins that fell into the top tenth percentile. **C)** APP (Aβ) and MAPT (Tau) protein levels. The central bar depicts mean and box edges indicate 25th and 75th percentiles, with whiskers extending to the 5th and 95th percentiles, excluding outlier measurements. **D)** Heat map representing the fold-change over the mean of mixed charge proteins in the top tenth percentile across the control and AD cases. Gene symbols are highlighted by their module color in U1-70K interactome (Black=not in a module). **E)** Gene ontology (GO) analysis of the 28-enriched mixed charge domain proteins highlights functions in RNA binding and processing. Significant over-representation of the ontology term is reflected with a Z>1.96, which is equivalent to p<0.05 (above red line).

### The LC1 domain of U1-70K interacts with pathological Tau from AD brain

In addition to aggregating in AD brain, spliceosomal proteins such as U1-70K associate with neurofibrillary tangles and co-localize with Tau paired helical filaments by electron microscopy (5–7,50). Furthermore, similar to the biophysical properties of RBPs, recent evidence now indicates that Tau undergoes LLPS *in vitro* (51). This process is enhanced by polyanions, such as heparin (51) and RNA (52), as well as phosphorylation on Tau (51). Based on these observations, we sought to assess if the LC1 domain of U1-70K could interact with pathological Tau from human AD brain. Equivalent amounts of GST purified LC1 domain or the N-terminal domain of rU1-70K were added to AD brain homogenates and immunoprecipitated with anti-myc antibodies followed by a western blot for Tau (**Fig. 9A**). Compared to the N-terminal domain, the LC1 domain of rU1-70K co-immunoprecipitated significantly more Tau, and modified Tau species of altered molecular weights (**Fig. 9B**). This indicates that the association of U1-70K and other mixed charge RBPs with Tau in AD are likely mediated by direct or indirect interactions via MC domains (**Fig. 10**). These findings highlight a shared role for MC domains in stabilizing protein-protein interactions and potentially mediating their co-aggregation with Tau in AD (**Fig. 10**).

**Figure 9.**
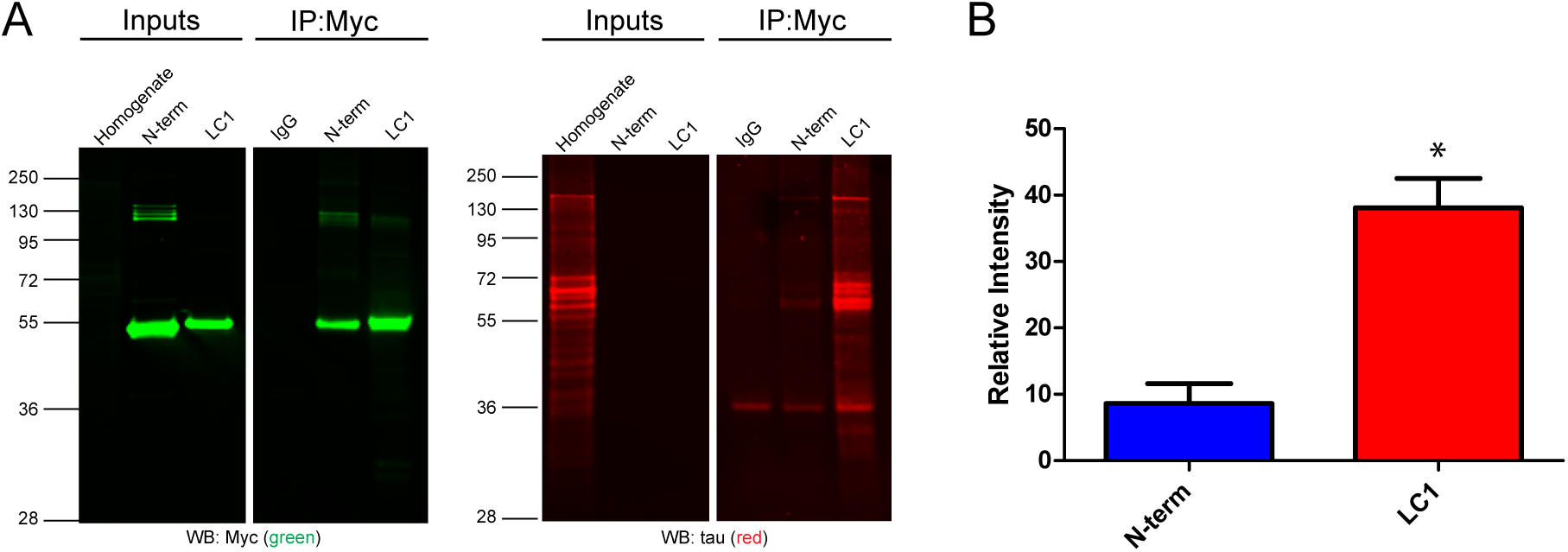
The mixed charge LC1 domain of U1-70K interacts with Tau in AD brain. **A)** Purified N-terminal or the mixed charge LC1 domain (4 µg) of rU1-70K were added separately to AD brain homogenates and immunoprecipitated with anti-Myc antibodies. IP with a non-specific IgG was also performed as a negative control. Inputs and immunoprecipitates were analyzed by Western blot using anti-Tau antibodies (red) and myc antibodies (green). **B)** The LC1 domain interacted with significantly higher levels of Tau from AD brain than the N-terminal domain (t-test one-tail p value = 0.0156). The experiment was done in biological triplicate (*n*=3) from independent AD cases (**Table 2)**.

**Figure 10.**
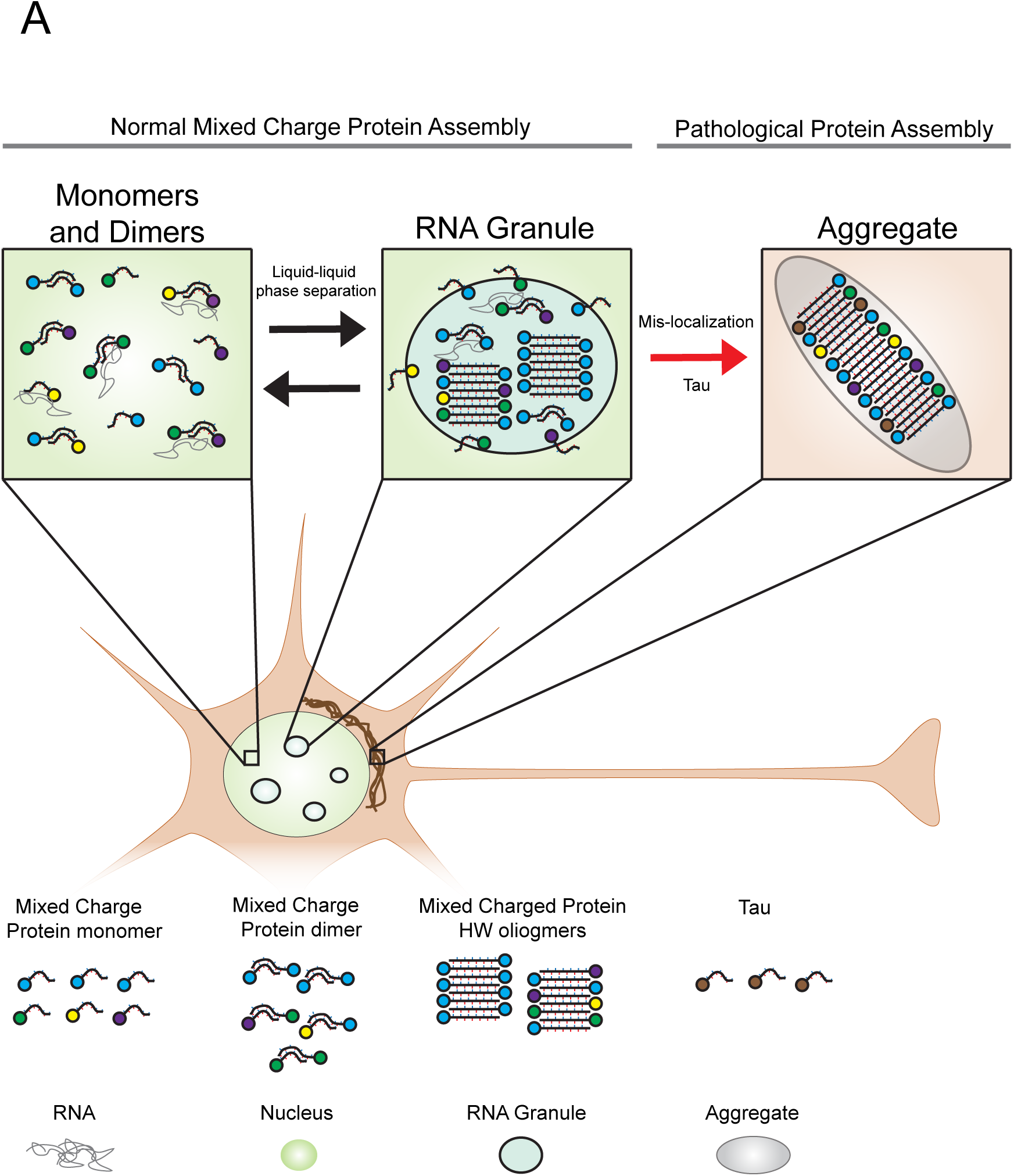
A Model for the assembly and pathological aggregation of U1-70K and other RNA-binding proteins with mixed charge domains in AD. RNA binding proteins with mixed charge domains reciprocally interact to form dimers, oligomers, and RNA granules under normal endogenous conditions. In AD, RNA binding proteins with mixed charge domains and tau co-aggregate in the cytoplasm to form aggregates under pathological conditions.

## DISCUSSION

In this study we show that the mixed charge LC1 domain of rU1-70K can directly self-interact *in vitro* to form high molecular weight oligomers and that this domain is also necessary and sufficient for U1-70K self-association in cells. Using quantitative proteomics, functional classes of U1-70K interacting proteins were identified that favored interactions with the LC1 domain. This revealed a class of structurally similar RBPs that also contained analogous mixed charge LC domains. We show that for at least two other RBPs, LUC7L3 and RBM25, their respective mixed charge domains are required for reciprocal interactions with U1-70K and for proper localization to nuclear granules. Global analysis of the detergent-insoluble proteome in human brain revealed elevated levels of mixed charge RBPs in AD. Finally, we show that the LC1 domain of U1-70K can interact with Tau from AD brain supporting a hypothesis that mixed charge structural motifs on U1-70K and related RBPs could mediate cooperative interactions with Tau in the pathology of AD.

We propose that mixed charge proteins be considered a class of proteins due to their related biological function, and shared primary structure. For example, we provide evidence to support the function of the LC1, and more broadly other mixed charge (MC) domains, in nuclear RNA granule formation. Thus, these reciprocal interactions of MC domains could form the “glue” that drives granule assembly. It has been previously proposed that MC domains, such as those found in U1-70K, RBM25 and LUC7L3, self-assemble through the formation of polar zippers (27). Furthermore, the promiscuous nature of MC domain interactions could be crucial to facilitating the complicated and multifaceted function of RNA processing proteins (16,53). The presence of distinct LC domains like the LC2 (317–407 residues) in U1-70K may further refine the dynamics of this process. For example, U1-70K interacts with both the spliceosome and members of the polyadenylation complex, including FIP1L1. Analogous to U1-70K, FIP1L1 harbors MC domain, and is the top hub protein in the blue module (54). Thus, the presence of MC domains in U1-70K enable physical, if not also functional, crosstalk between the role of U1-70K in 5’-splice site recognition and the polyadenylation complex in mRNA processing.

U1-70K is normally nuclear, but is found mislocalized to cytoplasmic Tau-immunoreactive neurofibrillary aggregates in AD neurons (6), which may contribute to a loss of spliceosome function given recently identified RNA splicing deficits in the disease (8). Mislocalization of other RBPs contributes to neurodegenerative disease (55,56). Here we show that the LC1 domain is important for nuclear localization of U1-70K, supporting a link between aberrant LC1 interactions and mislocalization of U1-70K. Our findings also shed light on previous studies in which the C-terminal domain (residues 161–437) of U1-70K was found sufficient for nuclear localization (30,57). Moreover, our ΔLC1+2 mutant protein mimic findings for the 1–199 U1-70K C-terminal truncation, wherein expression of this N-terminal fragment localized to the nucleus, but not to granules. It is likely that the nuclear localization sequence within the LC1 domain was missed by earlier studies due to the selected sites of truncation, as the LC1 domain was never expressed in its entirety (57).

We show that U1-70K and other mixed charge proteins share many of the same properties as Q/N-rich prion-like proteins, despite their difference in primary sequences. These properties include the ability to self-assemble into high molecular weight oligomers, form nuclear granules in cells, and promote aggregation. The formation of RNA granules has been viewed as an intermediary step towards protein aggregation (1,9,58) and our observations place U1-70K and other mixed charge proteins among prion-like RBPs, such as TDP-43, FUS, hnRNPA1, and TIA-1 that form granules and aggregate in neurodegenerative disease (4,21,26,59). Thus, the ability of the LC1 and other MC domains to self-interact poises them for pathological aggregation in neurodegenerative diseases, which is consistent with their increased insolubility in AD.

In AD brain, both U1-70K and Tau aggregate and co-localize to neurofibrillary tangles (5–7,50,60). However, the mechanisms underlying the relationship between Tau and U1-70K co-aggregation is unknown. Tau undergoes LLPS via electrostatic static interactions *in vitro*, referred to as coacervation, (52,61). Moreover, Tau LLPS is mediated by the Tau microtubule binding repeats (residues 244–369) (62). This is notable as a recent study examining the physical structure of Tau filaments in AD brain revealed an exposed mixed charge region within the tertiary structure of Tau, comprised of residues 338–358 in the microtubule binding repeat domain (63). Thus, it is tempting to speculate that pathological Tau in AD may behave like other mixed charge RBPs and sequester U1-70K to neurofibrillary tangles or vice versa. Specifically, we propose that this MC surface in pathological Tau mediates physical interactions between Tau and the U1-70K LC1 domain (**Fig. 10**). Furthermore, Tau-U1-70K hetero-oligomers may have a unique aggregation propensity, though additional determinants of aggregation may reside in the cytoplasm including RNA (50). Future studies examining these interactions in the context of the time-course of disease progression and with resolution of spatially defined interactions within neurons could illuminate the mechanisms of U1-70K aggregation with Tau.

In summary, we have identified novel functional roles for MC domains in protein-protein interactions, nuclear localization, granule formation, and pathological aggregation. We show similarities between MC domains and Q/N-rich domains found in RBPs that aggregate in neurodegenerative diseases. We also demonstrate how a weighted protein-protein interaction network analysis can be used to resolve biologically and structurally distinct complexes. Notably, RBPs with MC domains also demonstrated elevated insolubility in AD brain and the MC domain of U1-70K was sufficient to interact with pathological Tau. This supports a hypothesis that MC structural domains on U1-70K and other RBPs could mediate cooperative interactions with pathological Tau isoforms.

## EXPERIMENTAL PROCEDURES

### Materials

Primary antibodies used in these studies include an in-house rabbit polyclonal antibody raised against a synthetic KLH-conjugated peptide corresponding to a C-terminal epitope of U1-70K (EM439) (7), an anti-Myc-tag (clone 9B11, Cell Signaling), an anti-GST (ab6613 Abcam) antibody, an anti-LUC7L3 (HPA018484–100UL, Sigma), an anti-RBM25 (ab72237, Abcam), an anti-U1-70K monoclonal (05–1588, Millipore), an anti-tau (ab54193, Abcam), IgG mouse control (550339 BD Pharmigen). Secondary antibodies were conjugated to either Alexa Fluor 680 (Invitrogen) or IRDye800 (Rockland) fluorophores.

### Plasmids and Cloning

The original cDNA of U1-70K containing C-terminal myc and DDK tags was cloned from pCMV6-Entry vector (Origene) and inserted into the HindIII/BamHI sites in the pcDNA3.1 vector (5). Full-length and U1-70K deletions sequences were subsequently cloned into the EcoRV/XhoI sites in the pLEXM-GST vector for the expression of N-terminal and C-terminal GST-tagged proteins. Similar cloning strategies were performed using LUC7L3 (Origene RG214406) and RBM25 (Origene RC212256) plasmids. All cloning was performed by the Emory Custom Cloning Core Facility and plasmids confirmed by DNA sequencing.

### Immunoprecipitation

Human embryonic kidney (HEK) 293T cells (ATCC CRL-3216) were cultured in Dulbecco’s Modified Eagle Medium (DMEM, high glucose (Gibco)) supplemented with 10% (v/v) fetal bovine serum (Gibco) and penicillin-streptomycin (Gibco) and maintained at 37 °C under a humidified atmosphere of 5% (v/v) CO_2_ in air. For transient transfection, the cells were grown to 80–90% confluency in 10 cm^2^ culture dishes and transfected with 10 μg expression plasmid and 30 μg linear polyethylenimine. Cells were homogenized in ice-cold immunoprecipitation (IP) buffer containing (50 mM HEPES pH 7.4, 150 mM NaCl, 5% glycerol, 1 mM EDTA, 0.5% (v/v) NP-40, 0.5% (v/v) CHAPS, Halt phosphatase inhibitor cocktail (1:100, Thermo Fisher). Samples were sonicated for 5 seconds on 5 seconds off at 30% amplitude for a total of 1.5 minutes (13 cycles). The samples were cleared (14,000 x g for 10 minutes) and protein concentrations determined using a standard bicinchoninic acid (BCA) assay (Pierce). Protein A Sepharose 4B beads (Invitrogen 101042; 20 uL per IP), were washed twice in IP buffer and then blocked with 0.1 mg/ml bovine serum albumin (Thermo #23209) and washed three additional times in IP buffer. Then anti-Myc (4µg) mouse monoclonal antibody (Cell Signaling 2276) or 4 µg IgG control (550339 BD Pharmingen) was allowed to incubate rotating with the bead slurry in IP buffer (500µL) for a minimum of 90 minutes to allow antibody conjugation to beads. Beads were washed 3 times in IP buffer. Lysates where pre-cleared by centrifugation at 14,000 x g at 4^°^C for 10 minutes. The pre-cleared protein lysates were added to beads (1.5 mg per IP) and incubated rotating overnight at 4^°^C. The beads were washed 3 times in IP wash buffer (IP buffer without glycerol or CHAPS) by centrifugation at 500 x g for 5 min at 4^°^C then resuspended in IP wash buffer. Following the last wash, the bead suspension was transferred to a new Eppendorf tube to minimize contamination. The bound protein was eluted with 8M urea buffered in 10 mM Tris pH 8.0. For proteomics assays, 4 independent biological replicates were performed for each condition. For protein digestion, 50% of the eluted protein samples were reduced with 1 mM dithiothreitol (DTT) at 25°C for 30 minutes, followed by 5 mM iodoacetamide (IAA) at 25°C for 30 minutes in the dark. Protein was digested with 1:100 (w/w) lysyl endopeptidase (Wako) at 25°C for 2 hours and diluted with 50 mM NH_4_HCO_3_ to a final concentration of less than 2M urea. Samples were further digested overnight with 1:50 (w/w) trypsin (Promega) at 25°C. Resulting peptides were desalted with in-house stagetips and dried under vacuum.

### RNAase A Treatment

Cells were lysed in IP buffer with the addition of 5 mM MgCl_2_. Following sonication and centrifugation as described above, the lysates were split and treated with RNAase A or buffer alone (control) to a final concentration of 50 µg/ml of RNAase A. The RNAase A treated and control samples were incubated for 30 minutes at room temperature followed by centrifugation at 10,000 x g for 10 minutes at 4^°^C. The supernatant was added to beads and the IP was completed as detailed above.

### Blue Native Gel Electrophoresis

Recombinant N-terminal (residues 1–99) and LC U1-70K fragments (residues 231–308) were purified and their concentrations determined as described previously (5). The purified GST used as control was kindly gifted by the Dr. Richard Kahn (Emory University, Department of Biochemistry). Prior to analysis purified rU1-70K fragments were cleared by centrifugation at 20,000 x g for 15 minutes at 4^°^C to remove any insoluble precipitates. Each protein (0.8µg) was added to blue native gel loading buffer (5% glycerol, 50mM TCEP, 0.02% (w/v) G250 coomassie, 1x Native Page Running Buffer [Invitrogen BN2001]) and allowed to incubate at room temperature for 30 minutes. Samples were loaded onto a 3–12% NativePAGE Bis-Tris Gel (Invitrogen BN2011BX10) in addition to a native gel molecular weight marker (Thermo LC0725). Samples were resolved by electrophoresis at 150 volts for 1.5 hours in anode Native PAGE Running Buffer (Invitrogen BN2001) and cathode buffer with additive (Invitrogen BN2002). Gels were de-stained overnight in a solution of 15% (v/v) methanol and 5% (v/v) acetic acid and protein visualized on the Odyssey Infrared Imaging System (Li-Cor Biosciences). For western blot analysis, native gels were prepped with a 30 minutes incubation at room temperature in 1% (v/v) SDS and then transferred using the semidry iblot transfer system (Invitrogen) onto nitrocellulose (IB23001).

### Immunocytochemistry

Cells were plated on Matrigel (Corning #356234) coated coverslips and prepared for transfection using lipofectamine (ThermoFisher) according to manufacturer’s protocol. Immunocytochemistry was performed 48 to 72 hours after transfection essentially as described (64). After the blocking step, slides were dabbed to remove excess liquid and incubated in primary antibody overnight at 4^°^C. Primary antibodies included: rabbit anti-U1-70K (EM439), mouse anti-Myc-tag, mouse anti-LUC7L3, mouse anti-RBM25, mouse anti-U1-70K. The slides were washed 3 times with PBS 0.05% (v/v) saponin then incubated with secondary antibody (Dylight 549, Alexa 488) for one hour shaking at room temperature. Again, slides were washed 3 times with PBS with 0.05% saponin. DAPI diluted in PBS was added to each slide and incubated for at least 30 minutes rotating at room temperature. Following additional rinses in PBS, cells were mounted in Vectashield (Vector Laboratories, Burlingame, CA) and sealed with nail polish. Images were captured on an FLUOVIEW FV1000 confocal laser scanning microscope (Olympus).

### Liquid chromatography coupled to tandem mass spectrometry (LC-MS/MS)

Tryptic peptides were analyzed by LC-MS/MS essentially as described (60). Peptides were resuspended in loading buffer (0.1% formic acid, 0.03% trifluoroacetic acid, 1% acetonitrile) and separated on a self-packed C18 (1.9 um Dr. Maisch, Germany) fused silica column (20 cm x 75 μM internal diameter; New Objective, Woburn, MA) by a NanoAcquity UHPLC (Waters, Milford, FA) and monitored on a Q-Exactive Plus mass spectrometer (ThermoFisher Scientific, San Jose, CA). Elution was performed over a 140-minute gradient at a rate of 300 nL/min with buffer B ranging from 3% to 80% (buffer A: 0.1% formic acid and 5% DMSO in water, buffer B: 0.1 % formic and 5% DMSO in acetonitrile). The mass spectrometer cycle was programmed to collect one full MS scan followed by 10 data dependent MS/MS scans. The MS scans (300–1800 m/z range, 1,000,000 AGC, 100 ms maximum ion time) were collected at a resolution of 70,000 at m/z 200 in profile mode and the MS/MS spectra (2 m/z isolation width, 28 normalized collision energy (NCE), 50,000 AGC target, 50 ms maximum ion time) were acquired at a resolution of 17,500 at m/z 200. Dynamic exclusion was set to exclude previous sequenced precursor ions for 30 seconds. Precursor ions with +1, and +6 or higher charge states were excluded from sequencing. The mass spectrometry proteomics data have been deposited to the ProteomeXchange Consortium via the PRIDE partner repository with the dataset identifier PXD008260.

### Database Search

Raw data files were analyzed using MaxQuant v1.5.2.8 with Thermo Foundation 2.0 for RAW file reading capability (65). The search engine Andromeda was used to build and search a concatenated target-decoy UniProt Knowledgebase (UniProtKB) containing both Swiss-Prot and TrEMBL human reference protein sequences (90,411 target sequences downloaded April 21, 2015), plus 245 contaminant proteins included as a parameter for Andromeda search within MaxQuant (66). Methionine oxidation (+15.9949 Da), asparagine and glutamine deamidation (+0.9840 Da), and protein N-terminal acetylation (+42.0106 Da) were variable modifications (up to 5 allowed per peptide); cysteine was assigned a fixed carbamidomethyl modification (+57.0215 Da). Only fully tryptic peptides were considered with up to 2 miscleavages in the database search. A precursor mass tolerance of ±20 ppm was applied prior to mass accuracy calibration and ±4.5 ppm after internal MaxQuant calibration. Other search settings included a maximum peptide mass of 6,000 Da, a minimum peptide length of 6 residues, 0.05 Da tolerance for high resolution MS/MS scans. The false discovery rate (FDR) for peptide spectral matches, proteins, and site decoy fraction were all set to 1%. The label free quantitation (LFQ) algorithm in MaxQuant (33,67) was used for protein quantitation.

### Protein-protein interaction network analysis

The R package Weighted Gene Correlation Network Analysis (WGCNA) was used to sort proteins into functional groups by examining relative levels of co-enrichment (37). In WGCNA, correlation coefficients between each protein pair in the dataset are first calculated and transformed continuously with the power adjacency function to generate an adjacency matrix that defines the connection strength between protein pairs. This adjacency matrix is then used to calculate a topological matrix (TO), which measures the interconnectedness or correlation between two proteins and all other proteins in the matrix. All proteins are then hierarchically clustered (e.g. average linkage) using 1-TO as a distance measure and module assignments are subsequently determined by dynamic tree cutting (37). Threshold power Beta for reduction of false positive correlations (i.e. the beneficial effect of enforcing scale free topology) was sampled in increments of 0.5, and as the target scale free topology R² was approached, 0.1. The power selected was the lowest power at which scale free topology R² was approximately 0.80, or in the case of not reaching 0.80, the power at which a horizontal asymptote (plateau) was nearly approached before further increasing the power had a negative effect on scale free topology R². Other parameters were selected as previously optimized for protein abundance networks (60). Thus, for the signed network built on protein LFQ abundances obtained from IP-LC-MS/MS, parameters were input into the WGCNA::blockwiseModules() function as follows: Beta 10.9, mergeCutHeight 0.07, pamStage TRUE, pamRespectsDendro TRUE, reassignThreshold p<0.05, deepSplit 4, minModuleSize 15, corType bicor, and maxBlockSize greater than the total number of proteins. T-Distributed Stochastic Neighbor Embedding (tSNE) analysis was performed as described (68). Proteins with WGCNA intramodular kME>=0.50 were retained, and all duplicated values removed, as well as proteins with any missing values for the 16 non-IgG measurements. Then Rtsne R package Barnes-Hut-Stochastic Neighbor Embedding (SNE) Rtsne function was run on the LFQ expression matrix to reduce dimensionality from 16 to 2. The remaining points or proteins (*n*=375) were colored according to WGCNA module membership. Gene Ontology (GO) Elite analysis on each module was performed as described previously (60).

### Bioinformatic analysis of mixed charge proteins in the detergent insoluble proteome in AD brain

Quantitative proteomic analysis using isobaric tagging of sarkosyl-insoluble fractions (frontal cortex) from eight AD and six control cases was previously performed as described Supplementary proteomic data was downloaded from (49) and R was used to generate histograms, Fisher exact p-value, box plots, and the clustered heat map with the Nonnegative Matrix Factorization (NMF) package.

### Nuclear and Cytoplasmic Fractionation

The fractionation protocol was performed as essentially described in (69) with slight modifications. Briefly, after transfections with full-length rU1-70K plasmids or respective mutants, HEK cells were harvested by scraping and washed with PBS including 1x Protease Inhibitor Cocktail (Sigma). Cells were then spun down at 1000 x g at 4°C for 5 minutes and carefully resuspended in hypotonic buffer (10mM HEPES pH 7.9, 20mM KCl, 0.1mM EDTA, 1mM DTT, 5% glycerol, 0.5 mM PMSF and Halt phosphatase inhibitor cocktail [1:100, Thermo Fisher]) and incubated on ice for 15 mins. The detergent NP-40 was then added to a 0.1% (v/v) final concentration and cells were briefly agitated by vortex and left on ice for 5 additional minutes followed by centrifugation for 10 minutes at 4°C at 15,600 x g. affording the supernatant (S1) as the cytoplasmic fraction and the pellet (P1) as the nuclear fraction. To determine if difference in nuclear and cytoplasmic distribution were significant across conditions repeated measures ANOVA with post-hoc Tukey was performed in GraphPad Prism.

### Immunoprecipitation of rU1-70K fragments from human AD brain homogenate

Post-mortem frontal cortex tissue from pathologically confirmed AD cases were provided by the Emory Alzheimer’s Disease Research Center (ADRC) brain bank (**Supplemental Table 2)**. Neuropathological evaluation of amyloid plaque distribution was performed according to the Consortium to Establish a Registry for AD (CERAD) semi-quantitative scoring criteria (70), while neurofibrillary tangle pathology was assessed in accordance with the Braak staging system (71). Tissues were homogenized in NP-40 lysis buffer (25mM Tris-HCl (pH 7.5), 150mM NaCl, 1mM EDTA, 1% NP-40, 5% Glycerol + protease + phosphatase inhibitors) using a bullet blender (60) followed by centrifugation at 10,000 x g for 10 mins at 4^°^C to clear tissue debris. Immunoprecipitation was performed from 1 mg of brain homogenate from three independent AD cases. Homogenates were first pre-cleared using 30µL of Protein A-Sepharose conjugated beads (Invitrogen #101041) rotating at 4^°^C for one hour. GST purified rU1-70K fragments (4µg) were added independently to the pre-cleared homogenates. IP was performed using anti-Myc-tag (clone 9B11, Cell Signaling) in samples containing the rU1-70K proteins. An IgG control antibody (550339 BD Pharmigen) was used as negative control. Immunocomplexes were captured using Dynabeads Protein G magnetic beads (Invitrogen #1003D), which were washed 3 times using wash buffer (50mM Tris HCl, pH 8, 150 mM NaCl and 1% NP-40) followed by 5 min boiling in Laemmli sample buffer to elute bound proteins prior to western blot analysis.

### Western Blotting

Western blotting was performed according to standard procedures as reported previously (5,10,64). Samples in Laemmli sample buffer 8% glycerol, 2% SDS, 50mM tris pH 6.8, 3.25% β-mercaptoethanol were resolved by SDS-PAGE before an overnight wet transfer to 0.2 μm nitrocellulose membranes (Bio-Rad) or a semi-dry transfer using the iBlot2 system. Membranes were blocked with casein blocking buffer (Sigma B6429) and probed with primary antibodies (see materials) at a 1:1000 dilution overnight at 4 °C. Membranes were incubated with secondary antibodies conjugated to Alexa Fluor 680 (Invitrogen) or IRDye800 (Rockland) fluorophores for one hour at room temperature. Images were captured using an Odyssey Infrared Imaging System (Li-Cor Biosciences) and band intensities were quantified using Odyssey imaging software.

### Assessment of protein similarity to LC1 domain of U1-70K

The U1-70K LC1 protein similarity list was created using the Uniprot pblast feature (http://www.uniprot.org/blast/) using the following parameters: Target Database Human containing 160,363 entries (updated March 2015), E-threshold: 10, Matrix: Auto, Filtering: None, Gapped: Yes, Hits: 1000. The input blast entry was the LC1 domain of U1-70K (residues 231–308). The resulting list of proteins was subsequently filtered to remove un-reviewed entries, producing 255 proteins with E-values less than 0.005 and similarity to the LC1 domain of higher than 20 percent. The "biotin-isoxazole" list originates from a previous study (45) using the biotinylated isoxazole compound to precipitate proteins from HEK293 nuclear extracts. The "prion-like" list of RNA binding proteins originates from a previous work study using *in silico* methods to identify Q/N rich prion-like proteins (44). Finally, protein alignment was done using Clustal Omega multiple sequence alignment(https://www.ebi.ac.uk/Tools/msa/clustalo/).

## ACKNOWLEDGEMENTS

Support was provided by 5R01AG053960, the Accelerating Medicine Partnership for AD (U01AG046161), the Emory Alzheimer’s Disease Research Center (P50 AG025688), and the NINDS Emory Neuroscience Core (P30NS055077). N.T.S. is also supported in part by an Alzheimer’s Association (ALZ), Alzheimer’s Research UK (ARUK), The Michael J. Fox Foundation for Parkinson’s Research (MJFF), and the Weston Brain Institute Biomarkers Across Neurodegenerative Diseases Grant (11060). I.B was funded by a pre-doctoral T32 NINDS training grant 3T32NS007480–16 and F31 NRSA grant F31NS09385902. We also acknowledge Dr. Measho Abreha for technical advice on brain homogenization and immunoprecipitation conditions.

**Conflict of interest statement**. None declared.

**Author contributions:** Conceptualization, I.B. J.J.L., A.I.L., and N.T.S; Methodology, I.B., D.M.D., E.B.D. and N.T.S.; Investigation, I.B. D.M.D., and E.B.D; Formal Analysis, I.B., D.M.D., E.B.D. and N.T.S., Writing – Original Draft, I.B., and N.T.S,; Writing – Review & Editing, I.B., E.B.D, D.M.D., M.G., J.J.L., A.I.L. and N.T.S.; Funding Acquisition, I.B., A.I.L. and N.T.S.; Resources, M.G., A.I.L. and N.T.S.; Supervision, N.T.S.

## Supplementary Figures

**Supplemental Figure 1. Cluster dendrogram of rU1-70K interacting proteins quantified mass spectrometry analysis.** The WGCNA cluster dendrogram stratified rU1-70K interacting proteins (*n* = 716) into distinct modules (M1-7) defined by dendrogram branch cutting. The correlation of each protein to the specific traits (LC1, LC2, and IgG) are portrayed in the heat map.

